# GenEPi: Piezo1-based fluorescent reporter for visualizing mechanical stimuli with high spatiotemporal resolution

**DOI:** 10.1101/702423

**Authors:** Sine Yaganoglu, Nordine Helassa, Benjamin M. Gaub, Maaike Welling, Jian Shi, Daniel J. Müller, Katalin Török, Periklis Pantazis

**Author notes:** Correspondence to: Periklis Pantazis.

## Abstract

Mechanosensing is a ubiquitous process to translate external mechanical stimuli into biological responses during development, homeostasis, and disease. However, non-invasive investigation of cellular mechanosensing in complex and intact live tissue remains challenging. Here, we developed GenEPi, a genetically-encoded fluorescent intensiometric reporter for mechanical stimuli based on Piezo1, an essential mechanosensitive ion channel found in vertebrates. We show that GenEPi has high specificity and spatiotemporal resolution for Piezo1-dependent mechanical stimuli, exemplified by resolving repetitive mechanical stimuli of spontaneously contracting cardiomyocytes within microtissues, in a non-invasive manner.

## Main Text

Throughout an organism’s lifetime, cell mechanosensation (i.e. the ability to perceive and respond to mechanical stimuli in the form of shear stress, tension or compression) is essential in a myriad of developmental, physiological, and pathophysiological processes including embryogenesis, homeostasis, metastasis, and wound healing^1^. How these processes incorporate active feedback via force sensing at the cellular level is an area of active study, and, in recent years, a wide range of tools have been developed to interrogate cell mechanics^2,3^.

For instance, atomic force microscopy (AFM) and micropipette aspiration have proven to be powerful techniques to quantitatively measure tension in embryos and dissociated cells^4,5^. Other methods, which do not require direct and constant access to the sample, such as droplet-based sensors^6^ or optical^7^ and magnetic^8^ tweezers can modulate probes from a distance and allow precise measurement of molecular to tissue-level forces^2^. Still, these approaches typically require dissociated tissue or their use is complicated by the probe injection and size, which can damage the tissue^2^.

The necessity to non-invasively measure molecular forces in cells led to the development of genetically-encoded, Förster resonance energy transfer (FRET)-based fluorescent tension sensors, capable of measuring mechanical forces across specific cytoskeletal and adhesion proteins such as vinculin^9^, β-spectrin^10^ or cadherins^11^. As the specificity and force sensitivity of these probes is defined by the choice of protein and the FRET tension module, their use is restricted to a limited range of biological contexts and force regimes^2,3^.

Meanwhile, stretch-activated ion channels, including the Piezo proteins, are capable of responding to various external mechanical stimuli^12,13^. Most vertebrates have two Piezo genes — *Piezo1* and *Piezo2^12^*. While Piezo2 function is mainly restricted to the peripheral nervous system, Piezo1 is expressed in a wide range of tissues and has been shown to contribute to mechanotransduction in various organs^13^ (**Supplementary Table 1**). Mutations in human Piezo1 have been implicated in diseases such as dehydrated hereditary stomacytosis^14,15^ and general lymphatic dysplasia^16,17^. Global knockout of Piezo1 in mice causes embryonic lethality^18,19^, highlighting the importance of this channel for development and homeostasis^13^. How cells and tissues integrate Piezo1 activity has been mainly examined by outputs such as morphological changes, protein expression, electrophysiological signaling, cytosolic calcium (Ca^2+^) imaging, and transcriptional activity in response to mechanical stimuli^20^.

In order to develop a non-invasive, genetically-encoded fluorescent reporter for mechanical stimuli that is applicable to a wide variety of cells and types of mechanical stimuli, we set out to generate a reporter of Piezo1 activity. It has been recently shown that the C-terminus of Piezo1 resides within the cytosol and contains the ion-permeating channel^21,22^, which has a preference for divalent cations such as Ca^2+21,22^. Upon opening, Ca^2+^ concentration near the channel, referred to as Ca^2+^ microdomain, is typically several fold higher than resting levels^23^. We therefore hypothesized that by targeting a genetically-encoded Ca^2+^ indicator (GECI) to the ion permeating channel of Piezo1, we can obtain an optical readout for its activity.

We reasoned that a fluorescent reporter of channel activation would require a GECI with low Ca^2+^ affinity and a wide dynamic range to reliably monitor the considerable Ca^2+^ increase in the microdomains, while displaying a low response to cytosolic Ca^2+^, which serves as an important secondary messenger in many other cellular processes^23^. To meet these requirements, we decided to evaluate GCaMPs, a class of GECIs^24^, as fluorescent reporters of Piezo1 function. In contrast to FRET-based GECIs, GCaMPs occupy a narrow spectral range, allowing for the simultaneous imaging of multiple fluorescent markers. Progressive protein engineering efforts have yielded GCaMP variants that display a wide dynamic range of response with high signal-to-noise ratios (SNR)^25^.

In a systematic screen, we generated a library of reporters by fusing five different low-affinity GCaMPs^26,27^ (here denoted as GCaMP-G1 - GCaMP-G5) (with *K*_d_-s in the 0.6 to 6 µM range) to the C-terminus of human Piezo1 (Fig. 1a). Given the influence of linker length on the sensing mechanism^28^, we employed flexible linker peptides with varying lengths to attach GCaMPs to Piezo1 (**Supplementary Fig. 1a**). The generated variants were evaluated based on their response to both mechanical stimuli and cytosolic Ca^2+^ fluctuations that were independent of Piezo1 activity. To test their responses to mechanical stimuli, variants were exposed to physiological levels of fluid shear stress^29^ (see **Online Methods**) (**Supplementary Fig. 1b**), which causes a Piezo1-dependent Ca^2+^ increase in HEK 293T cells^18^. To determine the sensitivity of the variants to intracellular Ca^2+^ levels independent of Piezo1 function, we recorded their response to the Ca^2+^ ionophore ionomycin (**Supplementary Fig. 1b**)^30^.

**Figure 1.**
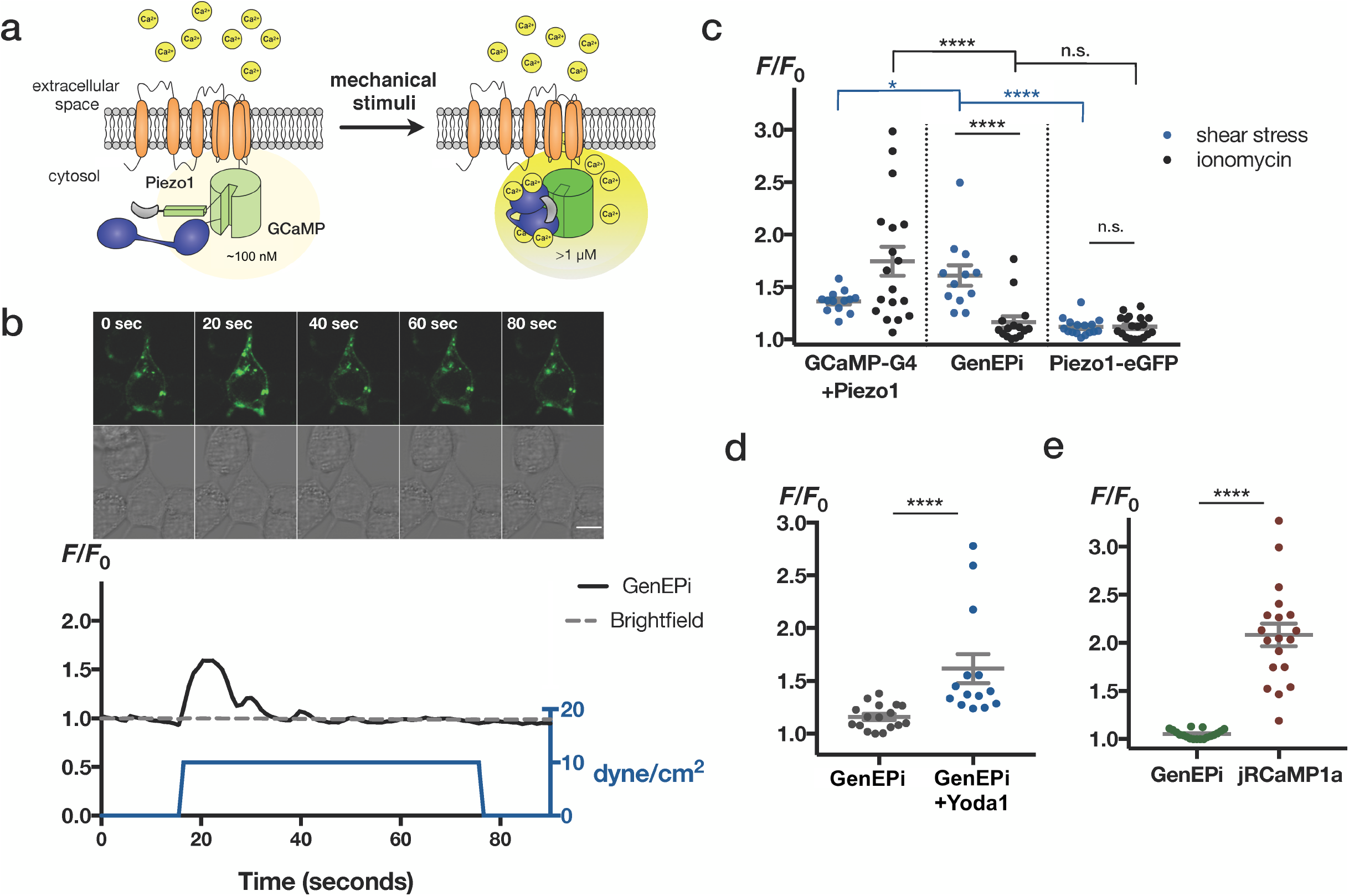
*In vitro* characterization of GenEPi shows that the reporter has functional specificity. (**a**) GenEPi sensing mechanism. GCaMP is targeted near the C-terminal Piezo1 channel. When mechanical stimuli induce channel opening, incoming Ca^2+^ (in yellow) binds to GCaMP, causing an increase in green fluorescence. (**b**) Representative example of GenEPi activation and *F/F*_0_ signal intensity profile (black) in response to 10 dyn/cm^2^ shear stress (blue) in HEK 293T cells. Time stamps in the images correspond to the stimulation and response profile in the graph. Scale bar, 10 μm. (**c**) Response of HEK 293T cells expressing Piezo1 and GCaMP-G4 (n=13, shear stress; n=18, ionomycin), GenEPi (n=12, shear stress; n=15, ionomycin), or Piezo1-eGFP (n=16, shear stress, n=19, ionomycin) to shear stress and ionomycin. Two-tailed Mann-Whitney test, ****=p<0.0001; *=p<0.05; n.s.=p>0.05, data from three independent experiments. (**d**) Response of GenEPi expressing HEK 293T cells to 10 μM Yoda1 (n=14) or DMSO (n=17). Two-tailed Mann-Whitney test, ****= p<0.0001, data from three independent experiments. (**e**) Response of GenEPi and jRCaMP1a expressing HEK 293T cells (dots, n=19) to intracellular Ca^2+^ triggered by 30 μM ATP. Welch’s t-test, ****= p<0.0001, data from six independent experiments. (**c, e**) Grey bars are means ±s.e.m.

Among the candidates tested, we identified one GCaMP-Piezo1 fusion variant that satisfied our requirements, Piezo1-1xGSGG-GCaMP-G4 (containing the GCaMP6s RS-1 EF-4 variant^27^), hereby referred to as GenEPi (Fig. 1c). GenEPi did not affect the viability of HEK 293T cells (**Supplementary Fig. 2**) and its localization in plasma membrane and endoplasmic reticulum reflected that of wild type Piezo1 (**Supplementary note 1** and **Supplementary Fig. 3**). The optical response of GenEPi (**Supplementary Note 1)** to fluid shear stress (Fig. 1b,c) was considerably higher (1.61 ±0.09, mean ±s.e.m, n=12 cells) than that of GCaMP6s RS-1 EF-4 (denoted here as GCaMP-G4) expressed in the cytosol (1.36 ±0.02, n=13 cells) (Fig. 1c), indicating that channel tethering of GCaMP-G4 in this particular configuration provides optimal access to high Ca^2+^ levels upon Piezo1 channel opening in response to mechanical stimuli. Importantly, cytosolic GCaMP-G4 could not distinguish between shear stress and ionomycin and responded to both stimuli (Fig. 1c). Furthermore, as GenEPi retained the low affinity for Ca^2+^ (**Supplementary Fig. 4a,b**), it had a low level of response to cytosolic Ca^2+^ induced by ionomycin (1.16 ±0.05, n=15 cells), indistinguishable from the response levels of the control fusion protein, Piezo1-eGFP (1.12 ±0.02, n=19 cells) (Fig. 1c). Interestingly, changing the level (1-30 dyn/cm^2^) or duration (10-120 sec) of fluid shear stress did not result in any significant difference in GenEPi’s response (**Supplementary Fig. 5a,b**), suggesting that the GCaMP response to high Ca^2+^ influx at the channel opening is not concentration-dependent, which confirms previous analysis demonstrating that the GCaMP-G4 response to Ca^2+^ binding is not linear but highly cooperative^27^. The functional specificity of GenEPi was validated by its selective response to the Piezo1-specific small molecule agonist Yoda1^31^, which significantly increased the reporter response (Fig. 1d).

In addition, we determined GenEPi’s response to physiological Ca^2+^ signaling in the cell upon addition of 30 µM ATP. We detected an ATP-dependent cytosolic Ca^2+^ increase using the Ca^2+^ indicator jRCaMP1a (Fig. 1e, **Supplementary Fig. 6**)^32^ and found that the elevated Ca^2+^ levels were detected by jRCaMP1a, however, not by GenEPi (**Supplementary Fig. 6**). These results indicate that GenEPi is indeed responding specifically to Piezo1-dependent activity and does not sense physiological fluctuations of cytosolic Ca^2+^, whereas cytosolic Ca^2+^ indicators respond to both Piezo1-dependent and Piezo1–independent stimuli (Fig.1c, Fig.1e, **Supplementary Fig.6**). The specificity of GenEPi’s response was further corroborated by the observation that membrane localization of GCaMP-G4 was not sufficient to confer functional specificity (**Supplementary Note 2**). Furthermore, channel tethering of all investigated GCaMP variants consistently reduced their response to cytosolic Ca^2+^ evoked by ionomycin (**Supplementary Fig. 1b**), which suggests that genetically-encoded Ca^2+^ indicators placed near the channel are protected from cytosolic Ca^2+^ fluctuations, supporting the microdomain hypothesis^23^. Taken together, GenEPi manifests high SNR and, in contrast to cytosolic Ca^2+^ indicators, demonstrates functional selectivity to the Piezo1-dependent fluid shear stress stimulus.

As Piezo1 is known to respond to other forms of mechanical stimuli, such as compression, we characterized the force sensitivity and temporal kinetics of GenEPi under this stimulus. We turned to a previously described AFM-based setup^33^ that allows probing Piezo1 sensitivity to mechanical stimuli while simultaneously recording the optical response of GenEPi (Fig. 2a,b). We applied precisely-timed compressive forces ranging from 100 nN to 400 nN with 50 nN increments on single HEK 293T cells expressing GenEPi using a 5 µm bead attached to an AFM cantilever (Fig. 2c). These compressive forces related to pressures ranging from 2.6 to 10.2 kPa or 19.1 to 76.5 mmHg (**Supplementary Note 3**). GenEPi responded to short (250 ms) compressive forces with fast kinetics, but on average with comparable signal amplitude to shear stress (1.65 ±0.12, n=21 cells) (Fig. 2d). While GenEPi signals in response to compressive forces were abolished in response to GsMTx-4, an inhibitor of Piezo1^34^ (Fig. 2e), the Piezo1-eGFP fusion did not show any optical response (Fig. 2d). The precise control of stimulation level and duration in this experimental setup allowed us to characterize the force sensitivity and duration of Piezo1-induced fluorescent signals reported by GenEPi and cytosolic GCaMP-G4. Although HEK 293T cells express small amounts of Piezo1, channel overexpression is required to reliably confer mechanical sensitivity to HEK 293T cells^35^ (**Supplementary Fig. 7**). To compare Piezo1-induced fluorescent signals, we applied timed compression onto GenEPi transfected cells and control cells co-transfected with human Piezo1 and cytosolic GCaMP-G4. Measured threshold forces were comparable for GenEPi and cytosolic GCaMP4 (243.50 ±13.68 nN and 241.20 ±13.87 nN, each n=21 cells, respectively) (Fig. 2f), demonstrating that the mechanical sensitivity of the channel is not affected by the protein fusion. Similarly, electrochemical response, ion selectivity and channel kinetics of Piezo1 within GenEPi were preserved in response to mechanical stimulation, the agonist Yoda1, and the generic inhibitor ruthenium red (**Supplementary Fig. 8-10**).

**Figure 2.**
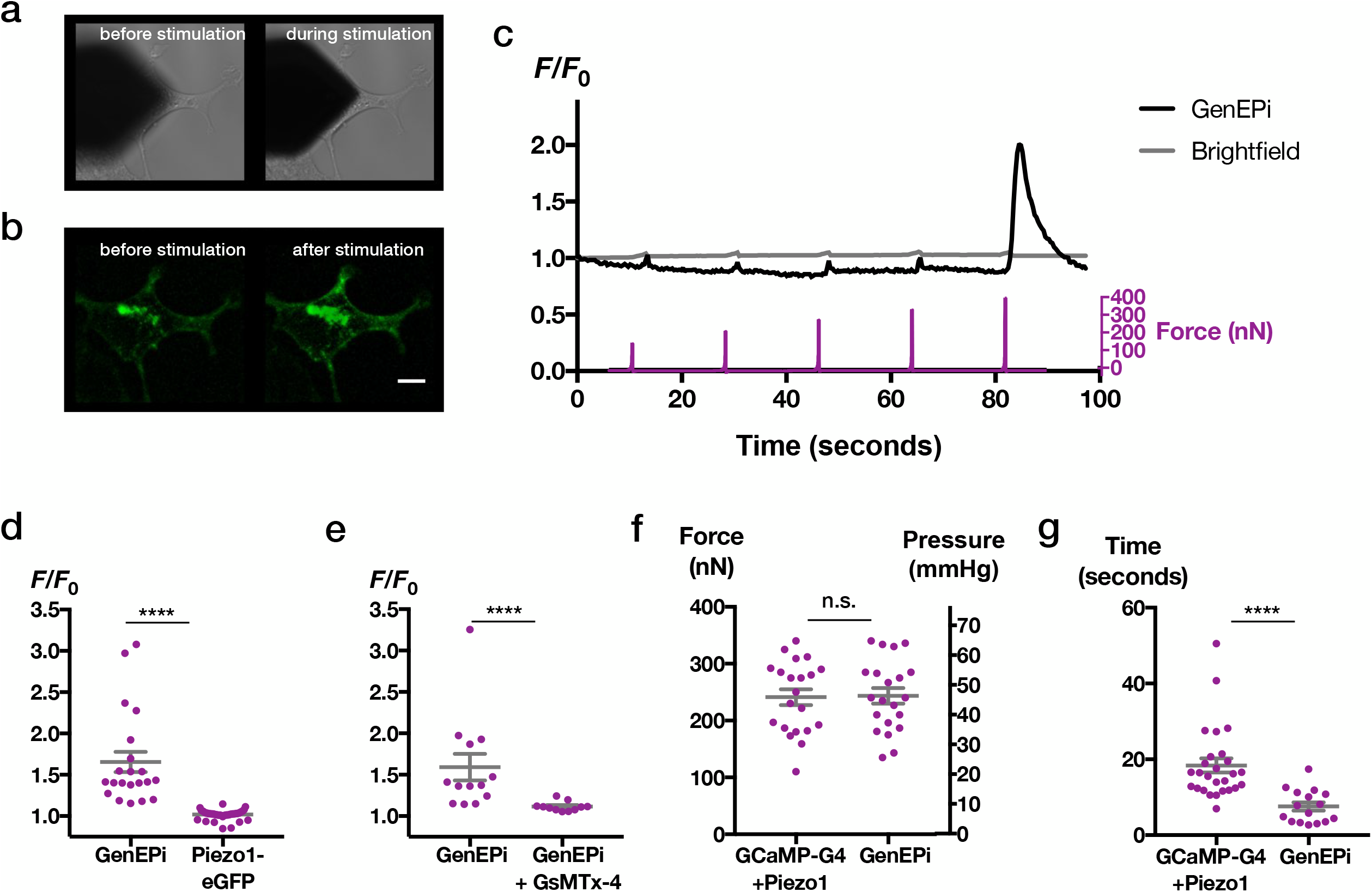
GenEPi retains force sensitivity and shows fast temporal resolution in response to compression. (**a**) Representative images of AFM cantilever stimulation of GenEPi expressing HEK 293T cells stimulated by the compressing AFM cantilever. Brightfield image of cantilever position before and during stimulation and (**b**) fluorescent image of the stimulated cell before and after stimulation. Scale bar, 10 μm. (**c**) The mechanical stimulation procedure, of compressive forces ranging from 129-370 nN (purple) along with the brightfield (grey) and fluorescent (black) traces from the cell depicted in **a,b**. (**d**) Amplitude of Ca^2+^ responses from GenEPi (n=21), and Piezo-eGFP (n=45) expressing cells. Two-tailed Mann-Whitney test, ****= p<0.0001. (**e**) Amplitude of Ca^2+^ responses from GenEPi expressing cells before (n=13) and after addition of 3 μM GsMTx-4 (n=11). Two-tailed Mann-Whitney test, ****= p<0.0001. (**f**) Threshold forces and pressures for cells co-transfected with human Piezo1 and cytosolic GCaMP-G4 (n=21), n.s.=p>0.05, Unpaired t-test. (**g**) Duration of Ca^2+^ responses from cells co-transfected with cytosolic GCaMP-G4 and human Piezo1 (n=27) and GenEPi (n=16). Two-tailed Mann-Whitney test, ****= p<0.0001. Grey bars are means ±s.e.m., data from three independent experiments.

GenEPi’s response to cantilever-triggered compression lasted on average 7.56 ±1.09 seconds (n=16 cells), which was much shorter than that of the cytosolic indicator (18.39 ±1.84 seconds, n=27 cells) (Fig. 2g), while the electrochemical inactivation kinetic of GenEPi in response to mechanical stimuli was comparable to Piezo1 and shorter than that of the Piezo1 delayed inactivation mutant R2456H^36^ (**Supplementary Fig. 11**). In conclusion, GenEPi provides not only high spatial resolution and functional specificity, when compared to cytosolic Ca^2+^ indicators, but also offers a gain in temporal resolution in response to mechanical stimuli.

In order to test GenEPi’s functional specificity and performance in a three-dimensional and multicellular environment, we tested its response to homeostatic cell motions, such as cardiomyocyte contraction. To this end, we generated doxycycline-inducible GenEPi mouse embryonic stem cells (mESCs)^37^ (**Supplementary Fig. 12a**) and differentiated these cells to cardiomyocytes^38^. We confirmed GenEPi’s activity in undifferentiated mESC by monitoring its specific response to Yoda1 (**Supplementary Fig. 12b**). After 10 days of differentiation (**Supplementary Fig. 13a)**, spontaneously beating patches of cells could be identified in microtissues (**Supplementary Video 1**) consisting predominantly of cardiomyocytes (**Supplementary Fig. 13d)**, and other mesodermal lineage cells, such as smooth muscle cells and endothelial cells (**Supplementary Fig. 13c,e**). Within some beating patches, we observed cells that displayed noticeable GenEPi responses to the cardiomyocyte contraction-triggered mechanical stimulation. The response amplitude range (*F/F*_0_ =1.15 to 2.29) was comparable to that of shear stress and compressive forces, yet the responses lasted less than a second (Fig. 3a-c and **Supplementary Video 2**). The subcellular and subsecond GenEPi response (Fig. 3a-c) rate was qualitatively coupled to the autonomous beating of the cardiomyocytes but at a slower frequency (**Supplementary Video 2**), which could be also observed in the electrochemical response of Piezo1 within GenEPi in response to repetitive mechanical stimulation (**Supplementary Fig. 14**).

**Figure 3.**
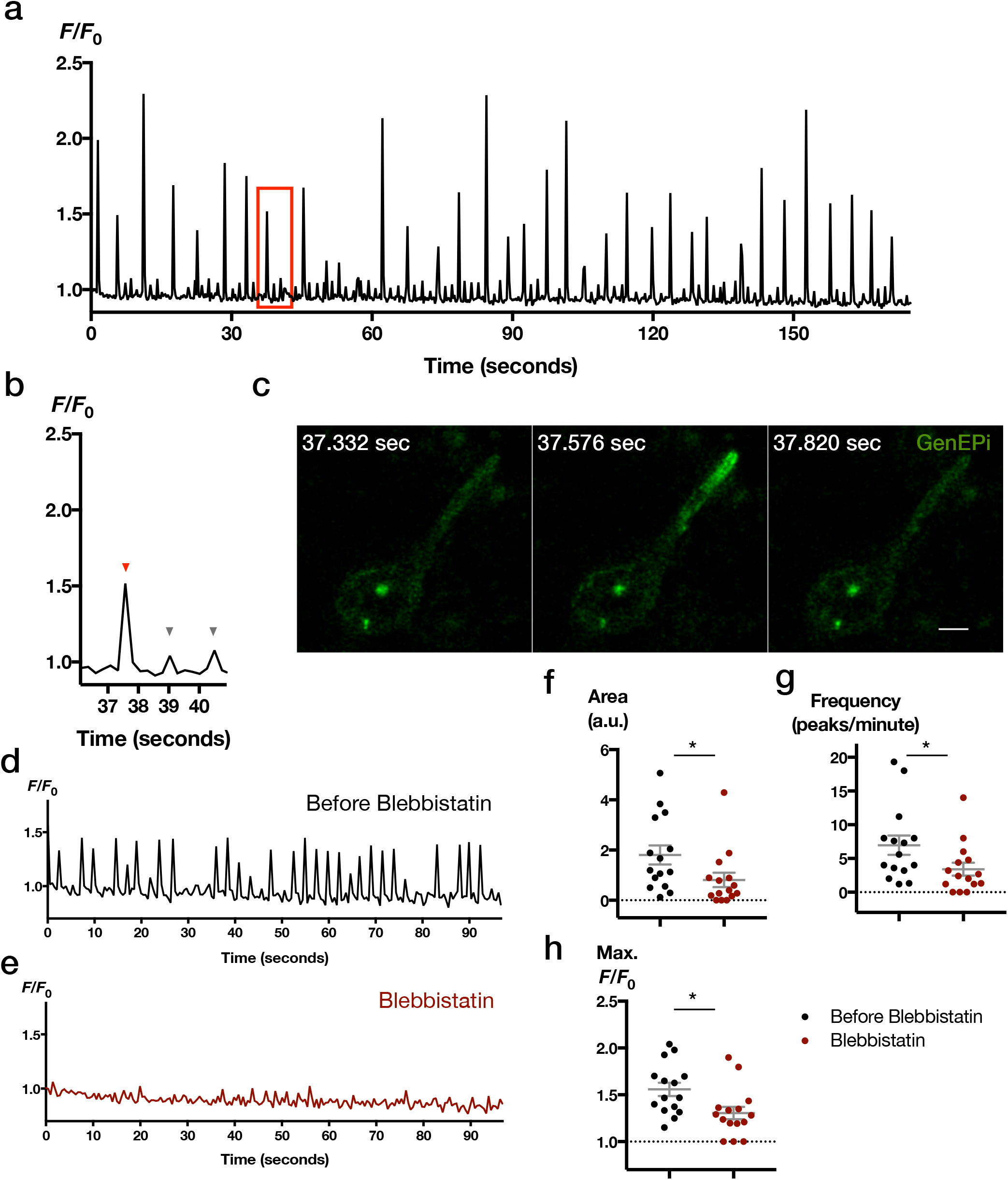
GenEPi reports cardiomyocyte contraction-triggered mechanical stimulation with high spatiotemporal resolution. (**a**) Intensity profile of a single cell attached to the beating patch expressing GenEPi in response to the autonomous beating of the cardiomyocytes. (**b**) Magnified intensity profile from the boxed region in red in (**a**) of GenEPi’s responses (red arrowhead) and fluorescence artifacts (grey arrowhead) upon cardiomyocyte contraction. (**c**) Time-lapse images of GenEPi response depicted in (**b**). Scale bar, 5 μm. (**d**) Representative *F/F*_0_ signal from a ROI in a cell in response to cardiomyocyte contraction. (**e**) Same ROI as in (**d**), after addition of 100 µM blebbistatin. (**f**) Total area, (**g**) frequency of peaks, and (**h**) maximum *F/F*_0_ measurement for each ROI before, and after the addition of blebbistatin. n=15 ROIs, error bar=s.e.m. Wilcoxon rank-sum test, *= p<0.05, **= p<0.01. Grey bars are means ±s.e.m., data from three independent experiments.

In order to confirm our observation that the source of these GenEPi responses were indeed cardiomyocyte contractions, we applied blebbistatin, a myosin inhibitor, which blocks the contractions and uncouples mechanical stimuli-induced Ca^2+^ influx from Ca^2+^ processes accompanying spontaneous cell contractions^39^. Fast GenEPi responses (Fig. 3d) decreased when contractions stopped in response to blebbistatin (Fig. 3e), as demonstrated by the significant decrease in the amplitude and frequency of the GenEPi response (Fig. 3f-h), confirming the cardiomyocyte contractions as the source of GenEPi signals. Hence, GenEPi shows a high spatiotemporal resolution of Piezo1 activity in microtissues, capable of specifically sensing repetitive and spontaneous mechanical stimuli of beating cardiomyocytes.

In summary, we introduced GenEPi as an intensiometric, genetically-encoded reporter for mechanical stimuli. GenEPi provides a specific and non-invasive functional readout of Piezo1 activity in response to mechanical stimuli, including shear stress and compressive forces with high spatiotemporal resolution in cells as well as small microtissues. This was achieved by successfully targeting a low-affinity GCaMP to the Ca^2+^ microdomain near the Piezo1 channel, which resulted in specificity for only Piezo1-dependent Ca^2+^ signals. Due to the highly cooperative Ca^2+^ sensing mechanism of GCaMP, GenEPi does not quantitatively report mechanical stimuli; however, GenEPi has a significantly broader applicability as compared to other genetically-encoded mechanical reporters, since Piezo1 has been identified to play a central role for mechanosensation in an increasing number of cell types and contexts (**Supplementary Table 1**). For instance, Piezo1 has been shown to sense mechanical properties of the environment of neural progenitor cells, influencing neuronal differentiation^40^, and to play a role in regulating volume of red blood cells^41^, where changes in shear stress and other mechanical forces are common. While the use of GenEPi requires overexpression of the Piezo1 channel; the role of GenEPi in various contexts can be studied using the inducible GenEPi mESC line, with full control over the expression levels of Piezo1, using doxycycline. Altogether, these features render GenEPi an ideal tool to elucidate the full extent to which mechanical signals, and more specifically Piezo1 channels, regulate development, physiology, and disease.

## ACKNOWLEDGEMENTS

We thank members of the Pantazis group, especially A.Y. Sonay for discussion and feedback. We thank W.P. Dempsey for help in designing the sketch depicting the mechanism of the reporter and feedback on the manuscript. This work was supported by the Swiss National Science Foundation (SNF grant no. 31003A_144048 to P.P.), the European Union Seventh Framework Program (Marie Curie Career Integration Grant (CIG) no. 334552 to P.P.), the Royal Society Wolfson Research Merit Award holder to P.P., the Rubicon grant from the Netherlands Organisation for Scientific Research and a fellowship from the Peter und Traudl Engelhorn Stiftung to M.W., the European Molecular Biology Organization (EMBO; ALTF 424-2016 to B.M.G.), the NCCR Molecular Systems Engineering, the Wellcome Trust Project Grant 094385/Z/10/Z and BBSRC Project Grant (BB/M02556X/1 to K.T.) and British Heart Foundation Intermediate Basic Science Research Fellowship (FS/17/56/32925 to N.H. and FS/17/2/32559 to J.S.). The Institute of Translational Medicine, Cellular and Molecular Physiology, University of Liverpool is thanked for the use of Zeiss LSM 800 confocal microscope.

## AUTHOR CONTRIBUTIONS

S.Y. conceived and S.Y. and P.P. refined the idea. S.Y. designed, carried out, and analyzed all experiments with the exception of the following: K.T. designed the four GCaMPs (G1-G4), N.H. carried out *in situ* affinity measurements; B.M.G. and D.J.M. designed and carried out AFM-based force spectroscopy and simultaneous confocal microscopy; M.W. generated the dox-inducible mESC line and carried out differentiation experiments; J.S. carried out patch-clamp electrophysiology. S.Y. and P.P. wrote the manuscript and all authors contributed to editing the manuscript. P.P. supervised the project.

## DATA AVAILABILITY STATEMENT

The data that support the findings of this study are available from the corresponding author upon request.

## ONLINE METHODS

### Molecular cloning

We obtained the human Piezo1 cDNA from Kazusa Inc, Japan. Generation of the first four types of GCaMPs; mGCaMP6s-EF4 (GCaMP-G1), mGCaMP6f-EF4 (GCaMP-G2), mGCaMP6s RS1-EF3 (GCaMP-G3) and mGCaMP6s RS1-EF4 (GCaMP-G4), were described elsewhere^1^. Fast-GCaMP-EF20 (here denoted as GCaMP-G5) was a gift from Samuel Wang (Addgene plasmid #52645)^2^. pGP-CMV-NES-jRCaMP1a was a gift from Douglas Kim (Addgene plasmid #61462)^3^.

Piezo1 was amplified using Herculase II fusion DNA Polymerase (600675, Agilent Technologies) and all Ca^2+^ indicators with various linker lengths were amplified with Phusion high-fidelity DNA polymerase (M0530S, NEB). List of primers, ordered from Sigma-Aldrich, can be found in **Supplementary Table 2**. Piezo1 and the Ca^2+^ indicators were introduced using restriction cloning and T4 DNA ligase (NEB). The Lck targeting sequence flanking restriction sites were synthesized by Genewiz, and introduced upstream of GCaMP-G4 and GCaMP-G5. All restriction enzymes were purchased from NEB. PCR and digestion products were purified using QIAquick PCR purification kit (28104, Qiagen) and QIAquick Gel Extraction kit (28704, Qiagen). Ligations were carried out using T4 Ligase (NEB) at 24°C for 1 hour followed by chemical transformation using Turbo ultracompetent *E.coli* based on K12 strain (NEB) and grown on Agar LB plates (Q60120 and Q61020, Thermo Fisher) and LB liquid media (244610, BD Bioscience) supplemented with appropriate antibiotics (100 μg ml^−1^ Ampicillin or 50 μg ml^−1^ Kanamycin, Sigma-Aldrich). Clones were screened using restriction digest and sequenced by Microsynth. Plasmid DNA isolation was carried out using ZR Plasmid Miniprep (D4054, Zymo Research).

### Cell culture and transfection

HEK 293T cells were obtained from ATCC (ATCC CRL-3216). Cells were cultured at 37°C, 5% CO_2_, in high glucose DMEM with GlutaMAX (10569010, Thermo Fisher), supplemented with 10% FBS (P40-37500, Pan Biotech) and 1X Penicillin-Streptomycin solution (15140122, Thermo Fisher). Cells were routinely tested and were negative for mycoplasma infection using Mycoplasma detection kit (B39032, LuBioScience GmBH). Plasmid DNA for transfection was isolated from 50 ml LB culture (244610, BD Bioscience) containing appropriate antibiotics using the Zymopure Plasmid Midiprep kit (D4200, Zymo Research). The amount of DNA was measured using the Nanodrop 2000c Spectrophotometer (Thermo Fisher) and 400-800 ng of each plasmid was introduced into cells using nucleofection (V4XC-2024, Lonza). Briefly, 70-80% confluent cells were washed with 1X Dulbecco’s Phosphate Buffered Saline (DPBS) (D8537-500ML, Sigma-Aldrich) and dissociated using 0.05% Trypsin-EDTA (25300054, Thermo Fisher). 10 μl of the cell suspension was mixed with 10 μl Trypan Blue (0.4%) (15250061, Thermo Fisher) and added to the cell counting slide (1450011, Bio Rad Laboratories AG). Cell count and cell viability were automatically calculated by the TC10 Cell Counter (Bio Rad Laboratories AG) and only cell suspensions with >90% viability were used for transfections. For fluid shear stress and chemical treatment experiments, 10^6^ cells were centrifuged for 5 minutes at 90xg and resuspended in 100 μl of SF cell line nucleofector solution. The cells mixed with 400-800 ng of each plasmid were then transferred into the nucleofection cuvette and pulsed using the program CM-150 (V4XC-2024, Lonza). 500 μl of fresh media was added to the cells, which were then transferred to a well in a 6-well plate (140675, Thermo Fisher). At 24 hours post transfection, the cells were dissociated and counted as previously described. The cells were then seeded onto ibitreat flow chambers (Ibidi u-slide-VI 0.4, 80606, Ibidi GmbH) for fluid shear stress experiments or ibitreat coated 8-well slides (80826, Ibidi GmbH) with a density of 75,000 cells per channel or well.

For experiments to test the response of the reporter to various chemicals, we used 1 μM ionomycin (I3909-1ML, Sigma-Aldrich), 30 μM ATP (A6559-25UMO, Sigma Aldrich) diluted in DPBS, 10 μM Yoda1 (5586, Tocris Bioscience) diluted in DMSO (D8418, Sigma Aldrich) or 2.5 μM GsMTx-4 (Pepta Nova GmBH) diluted in water.

For AFM experiments, HEK 293T cells were transfected using lipofection. Briefly, 1.5 10^6^ cells were seeded in a T25 flask (CLS430639, Sigma-Aldrich) the day before transfection. 4 µg of the reporter, or 1.6 µg of GCaMP-G4 and 3.6 µg of Piezo1 were diluted in 250 µl of Opti-MEM (31985062, Thermo Fisher), while 20 µl of Lipofectamine 2000 reagent (11668019, Thermo Fisher) was also diluted in 250 µl of Opti-MEM. After 5 minutes of incubation, the two solutions were mixed together and further incubated for 20 minutes. The solution was then introduced to cells and washed away with fresh medium after 4 hours. At 24 hours post transfection, the cells were dissociated and counted as previously described, and seeded onto 35 mm-wide cover-glass bottom Fluorodishes (FD35-100, World Precision Instruments), with a density of 300,000 cells per plate.

For the *in situ* affinity measurements, HeLa cells were cultured in DMEM containing non-essential amino-acids (Life Technologies), penicillin/streptomycin (100 U ml^−1^, 100 μg ml^−1^, respectively) and 10% heat inactivated FBS (Life Technologies) at 37°C in an atmosphere of 5% CO_2_. Cells were plated on 35 mm glass bottom culture dishes (MatTek) and allowed 24 hours to adhere before transfection with FuGENE HD (Promega) following the manufacturer’s instructions. Cells were maintained for 12-24 hours before being used in experiments.

### Generation of inducible GenEPi-mESC cell line (iGenEPi)

Doxycycline-inducible GenEPi-mESCs were generated using ZX1 mESCs carrying rtTA in the Rosa26 locus and dox-inducible cre flanked by self-incompatible LoxP sites in the HPRT locus^4^, kindly provided by Dr. Michael Kyba. ZX1 mESCs were cultured in DMEM (Life Technologies), 15% FBS (PAN Biotech), 2 mM L-Glutamine (Invitrogen), 1X non-essential amino acids, 0.1 mM β-mercaptoethanol, 100 U ml^−1^ leukemia inhibitory factor (Peprotech), 1 μM PD0325901 (Selleckchem) and 3 μM CHIR99201 (R&D Systems) on gelatin coated plates. Prior to electroporation, ZX1 mESCs were exposed to 500 nl ml^−1^ doxycycline for 24 hours. 1 × 10^6^ ZX1 mESCs were electroporated with 3 μg p2lox plasmid in which GenEPi was cloned between LoxP sites in a 0.4 cm electroporation cuvette at 230 mV, 500 μF and maximum resistance in a Biorad electroporator (Biorad Genepulser Xcell). 24 hours after electroporation, antibiotic selection was started with 300 μg/mL G418 (Sigma). Colonies that incorporated GenEPi were verified by FACS analysis and expanded. Dox-inducible GenEPi mESCs were differentiated to cardiomyocytes as previously described^5^. GenEPi mESCs were seeded as 500 cell / 20 μl in hanging drops on non-adherent plates to generate embryoid bodies (EBs) in EB medium, IMDM (Life Technologies), 20% FBS (PAN Biotech), 2 mM L-Glutamine (Invitrogen), 1X non-essential amino acids and 0.1 mM β-mercaptoethanol. After 2 days, EBs were transferred to uncoated petridishes. From day 3-5, 1 μM XAV939 was added to the culture conditions and EBs were plated on gelatin coated dished from day 4. Beating EBs appeared at day 10 of differentiation. Beating EBs were manually dissected and dissociated using 2 mg/ml Collagenase/Dispase (Sigma) to generate smaller beating patches and single cells (Supplementary Fig. 10). For blebbistatin experiments, 40-120 μM Blebbistatin (B0560, Sigma Aldrich) diluted in DMSO (D8418, Sigma Aldrich) was applied to the cells in 20 μM steps until contractions were stopped.

### Determination of cell viability and cell toxicity

Cell viability was determined using trypan blue exclusion assay. Briefly, cells in triplicates seeded in 6-well tissue culture plates (Thermo Fisher) were transfected with varying concentrations of GenEPi or GCaMP-G4 and human Piezo1. At 24 and 48 hours post transfection, cells were washed with 1X PBS twice and detached using 0.05% Trypsin-EDTA (25300054, Thermo Fisher). 10 µl of cell suspension was then mixed with 10 µl 0.4% Trypan Blue, and 10 µl of this mixture was added to the cell counting slide (C10228, Thermo Fisher) and measured using Countess II Automated cell counter (Thermo Fisher). The viability was expressed as a fold difference of the untreated samples for each time point.

To determine cell toxicity, lactate dehydrogenase (LDH) assay (Life Technologies) was used on GenEPi or GCaMP-G4 and human Piezo1 transfected cells according to manufacturer’s instructions. Briefly, GenEPi or GCaMP-G4 and human Piezo1 transfected cells in triplicates were seeded in 96-well tissue culture plates (167008, Thermo Fisher). After 48 hours, 10 µl of Cell Lysis buffer was added to a non-transfected cell triplicate and incubated for 45 minutes at 37°C, 5% CO_2_ to obtain maximum LDH activity. Afterwards 50 µl of each cell sample as well as the non-transfected cells for spontaneous LDH activity and maximum LDH activity was transferred into a new 96-well plate and mixed with 50 µl reaction mixture. Following 30 minutes of incubation at room temperature, 50 µl stop solution was added and the absorbance was measured at 490 nm and 680 nm using Tecan M1000 plate reader. To determine LDH activity, the absorbance values for 680 nm were subtracted from that of 490 nm. Percentage of cytotoxicity based on maximum LDH activity was determined as following: 100 x (cell sample LDH activity-spontaneous LDH activity)/(maximum LDH activity-spontaneous LDH activity).

### Fluid shear stress applications

We used the ibidi pump system (#10905, Ibidi GmBH). Fluid shear stress levels were calibrated and imaging solution viscosity of the perfusion solution was determined according to manufacturer’s instructions. Depending on the level of fluid shear stress applied, perfusion set yellow-green (#10964, for 5-30 dyn/cm^2^) or perfusion set white (#10963, for 1-5 dyn/cm^2^) were used. Representative fluid shear stress application traces are shown in Fig. 1c.

### Confocal microscopy

Images were acquired using the Zeiss 780 NLO Confocor 3 equipped with an argon laser for 458 and 488 nm excitation, a diode pumped solid-state laser for 561 nm excitation and a HeNe laser for 633 nm excitation. Images of single cells were acquired using the C Apo 40x/1.1 W DICIII objective, excited with 488 nm for reporter, and 561 nm for tdTomato and jRCaMP1a excitation, respectively. In order to ensure fast image acquisition, we imaged single cells in a small region of interest within the field of view, recording a single z-plane over several minutes. Live imaging of cells was carried out in Live Cell Imaging Solution (A14291DJ, ThermoFisher).

### Atomic force microscopy (AFM)-based force spectroscopy and simultaneous confocal microscopy

Prior to the experiment, 5 µm diameter silica beads (Kisker Biotech) were glued to the free end of tipless cantilevers (CSC-37, Micromash HQ) using UV glue (Dymax) and cured under UV light for 20 minutes. Cantilevers with beads were plasma treated for 5 minutes using a plasma cleaner (Harrick Plasma) to ensure a clean surface, and subsequently mounted on a standard glass cantilever holder (JPK Instruments) of the AFM. Cells cultured on glass-bottom Petri dishes were kept at 37ºC using a Petri dish heater (JPK Instruments). For the mechanical stimulation, an AFM (CellHesion 200, JPK Instruments) was mounted on an inverted confocal microscope (Observer Z1, LSM 700, Zeiss). Cantilevers were calibrated using the thermal noise method^6^. Mechanical stimulation protocols were programmed using the JPK CellHesion software. During the mechanical stimulus, the AFM lowered the bead on the cantilever onto the cell with a speed of 10 µm s^−1^ until reaching the preset force, kept the preset force constant for 250 milliseconds, and then retracted with a speed of 100 µm s^−1^. Preset forces were applied in intervals from 100 nN to 400 nN with 50 nN increments with the time between intervals ranging from 10–25 seconds. Representative mechanical stimulation traces are shown in Fig 2c.

Confocal imaging was performed using an inverted laser-scanning microscope (LSM 700, Zeiss) equipped with a 25x/0.8 LCI PlanApo water immersion objective (Zeiss). Time-lapse images were acquired with 100– 300 milliseconds time resolution and acquisition was initiated > 10 seconds before the onset of the mechanical stimulus. Time-lapse images of Ca^2+^ responses were analyzed using the built-in ZEN blue software.

### Patch-clamp electrophysiology

Human embryonic kidney 293T (HEK293T) cells were transfected with the plasmids using lipofectamine 2000 (Invitrogen). 48 hours after transfection, whole-cell and cell-attached patch-clamp recordings were made with the Axopatch-200B (Axon Instruments, Inc.) equipped with the Digidata 1550B and the pCLAMP 10.6 software (Molecular Devices, Sunnyvale, CA, USA) on the cells at room temperature. The tip resistance of recording glass pipettes was between 3 and 5 MΩ. The currents were sampled at 20 kHz and filtered at 2 kHz. The mechanical force was applied through a recording pipette using a Patchmaster-controlled pressure-clamp HSPC-1 device (ALA Scientific Instruments).

For whole-cell recordings, the external solution consisted of (in mM) 133 NaCl, 3 KCl, 2.5 CaCl_2_, 1 MgCl_2_, 10 HEPES and 10 glucose (pH 7.3 with NaOH). The pipette solution was composed of (in mM) 133 CsCl, 1 CaCl_2_, 1 MgCl_2_, 5 EGTA, 10 HEPES, 4 MgATP and 0.4 Na_2_GTP (pH 7.3 with CsOH).

For cell-attached recordings, the extracellular solutions were composed of (in mM) 140 KCl, 1 MgCl_2_, 10 glucose and 10 HEPES (pH 7.3 with KOH). The pipette solutions consisted of (in mM) 130mM NaCl, 5 KCl, 1 CaCl_2_, 1 MgCl_2_, 10 TEA-Cl and 10 HEPES (pH 7.3 with NaOH).

### *In situ* Ca^2+^ titration of GenEPi

GenEPi-transfected HeLa cells were permeabilized using 150 μM β-escin (in 20 mM Na^+^-HEPES, 140 mM KCl, 10 mM NaCl, 1 mM MgCl_2_, pH 7.2) for 4 minutes. The solution was replaced with “zero free Ca^2+^” solution (20 mM Na^+^-HEPES, 140 mM KCl, 10 mM NaCl, 1 mM MgCl_2_, 10 mM EGTA, pH 7.2) and various Ca^2+^ concentrations (0.001, 0.01, 0.1, 1, 10, 50, 500, 10000 µM free Ca^2+^) were applied in the presence of 10μM ionomycin and 4 μM thapsigargin. Free Ca^2+^ concentrations were calculated using the two-chelators Maxchelator program^7^.

Cells were examined with a Zeiss LSM 800 confocal microscope equipped with a 63x/1.4 Plan-Apochromat oil immersion objective and a 488 nm diode laser as excitation light source. Emitted light was collected through Variable Secondary Dichroics (VSDs) onto a GaAsP-PMT detector. The fluorescence signal was monitored over an elliptical region of interest (ROI) in the plasma membrane using the ImageJ program. Data obtained from 12 to 42 cells (from at least three independent experiments) was plotted and analysed on GraphPad Prism 6. The fluorescence dynamic range (*F*_max_-*F*_0_)/*F*_0_ or ∆*F*/*F*_0_ was expressed as mean±s.e.m. The Ca^2+^ dissociation constant (*K*_d_) and cooperativity (*n*) were obtained by fitting the data to the Hill equation.

### Live staining and immunochemistry

Live cell staining of cells was achieved using the Image-IT LIVE plasma membrane and nuclear labeling kit cell staining kit (I34406, Thermo-Fisher), and the ER tracker red (E34250, Thermo-Fisher) according to product specifications. For antibody stainings, cells were fixed in 4% paraformaldehyde (15714-S, Lucerna Chem AG) for 5 minutes, washed with PBS and blocked using Max Block blocking medium (15252, Active Motif) supplemented with 0.1% TritonX-100 (T8787, Sigma Aldrich). Cells were then incubated with anti-GFP antibody (ab6673, Abcam) or anti-Piezo1 antibody (ab82336, Abcam) diluted in Max Block blocking medium. After several washing steps with PBS, the cells were incubated with goat anti-rabbit Alexa Fluor-633 (A-21071) or donkey anti-goat Alexa Fluor-633 (A-21082, Thermo Fisher) as well as DAPI (62248, Thermo Fisher). Mouse embryos and mESCs were fixed in 10% paraformaldehyde (15714-S, Lucerna Chem AG) for 10 minutes, permeabilized with PBS supplemented with 0.1% TritonX-100 (T8787, Sigma Aldrich) and blocked with 10% Donkey Serum (17-000-121, Jackson ImmunoResearch) in PBS supplemented with 0.1% TritonX-100. Samples were then incubated with anti-GFP antibody (ab6673, Abcam), anti-mouse E-cadherin (AF748-SP, Techne AG), anti-cardiac Troponin T antibody (ab8295, Abcam), anti-smooth muscle myosin heavy chain II antibody (ab53219, Abcam) or anti-CD31/PECAM-1 antibody (AF3628, R&D systems) diluted in 10% Donkey Serum (17-000-121, Jackson ImmunoResearch) in PBS supplemented with 0.1% TritonX-100. After several washing steps with PBS, the samples were incubated with donkey anti-goat Alexa Fluor-594 (A-11058, Thermo Fisher), goat anti-mouse Alexa Fluor-568 (A-11004), goat anti-rabbit Alexa Fluor-633 (A-21071) or donkey anti-goat Alexa Fluor-633 (A-21082, Thermo Fisher) as well as DAPI (62248, Thermo Fisher).

### Image processing and analysis

For the shear stress experiments, and time-lapse upon experiments with application of a chemical, the cells were automatically segmented using a MATLAB script. Briefly, the signals from the cytosolic tdTomato or jRCaMP1a were automatically identified, and high intensity pixels were used to generate a mask. This mask was then applied to the time series images of the reporter and the cytosolic signal and single cell intensities were extracted for each time point. This information allowed us to get the traces for the intensiometric reporter response of each cell. Ca^2+^ responses were expressed as fluorescence levels normalized to baseline (*F/F*_0_). To obtain (*F/F*_0_), we divided the fluorescence levels (*F*) by the baseline fluorescence of the cell (fluorescence of the first five frames, *F*_*0*_).

During the AFM experiments, mechanical stimulation of cells with the cantilever caused cytosolic or membrane bound fluorophores to move in or out of the confocal imaging plane, creating fluorescence artifacts. These artifacts were clearly distinguishable from Piezo1 receptor mediated Ca^2+^ influx, since (i) they showed strong symmetry with stepwise increase and decrease of fluorescence (ii) were very short in duration, and (iii) appeared synchronously with the mechanical stimulus and were thus preceding the Ca^2+^ responses. Fig. 2c illustrates such an example of fluorescence artifacts and Ca^2+^ responses. Any fluorescent signal that was greater than the artifact was classified as “response”, anything below was defined as noise. The resulting signal trace was processed as described above. The duration of the signal was calculated by subtracting the first time point fluorescence signal is higher than artifact from the time point signal goes back to the baseline. Baseline was calculated as average fluorescence of 5 seconds preceding the stimulus.

### Statistical analysis

All data are expressed means ±s.e.m. Sample sizes (n) are provided in the text or figure legend of each experiment. Each experiment has been repeated independently at least 3 times. Each data set was subjected to Shapiro-Wilk normality test to determine whether the data set has a Gaussian distribution; p>0.05 indicated it has a Gaussian distribution, and p<0.05 indicated it did not. When all of the compared data sets had Gaussian distribution, two-tailed Student’s t-test was applied to compare two independent datasets; with an *F*-test to compare variances. When F-test resulted in p<0.05, Welch’s correction was applied to the t-test. When more than 2 datasets were present with Gaussian distribution, one-way ANOVA was used to compare datasets, followed by Holm-Sidak’s *post hoc* multiple comparisons test. When at least one of the compared data sets did not have a Gaussian distribution, Mann-Whitney test was applied to compare two independent datasets; and Wilcoxon rank-sum test was applied when the data sets were paired. When more than 2 datasets were present without Gaussian distribution, Kruskal-Wallis test was applied, followed by Dunn’s *post hoc* multiple comparisons test. For all statistics, p-value were reported, with n.s.=p>0.05, *=p<0.05, **=p<0.01, ***=p<0.001 and ****=p<0.0001.

